# Costs and benefits of using rhythmic rate codes

**DOI:** 10.1101/2021.04.24.441276

**Authors:** Erik J Peterson, Bradley Voytek

## Abstract

Neural oscillations are observed ubiquitously in the mammalian nervous system, and the benefits of oscillatory coding have been the topic of frequent analysis. Many prior studies focused on communication between populations which were already oscillating and sought to understand how these rhythms and overall communication interact. We take a contrary view here. In this paper, we focus on measuring the costs of translating from an aperiodic code to a rhythmic one. We study two models. The first is simulated independent populations of neurons subjected to a theta-band (6 Hz) pacemaker using Linear-Nonlinear Poisson (LNP) sampling. The second is a model of beta-gamma oscillations using biophysical neurons with self-organized dynamics. We measure benefits and costs in both models using information theory. In both models oscillations can only benefit communications by increasing spiking by specific amounts, in effect, correcting for “undersampling” of the stimulus. This is mechanistically similar to theories for how deep brain stimulation can enhance cognition and is consistent with older studies of gamma entrainment. Yet this trend was not universal. No one guiding principle of dynamics determines the cost of a translation in the models we studied. In our models to predict the benefits or costs of an oscillatory translation we need to understand the exacting physical details of the intrinsic connections, the population size, and the external drive.

---

In this paper we begin to reconcile the fact that oscillations are often found in the nervous system with the fact that many aspects of the natural world an animal must sense are not themselves oscillatory, and that whatever benefits adopting a oscillatory code are there may also be costs. In particular, we use information theory to measure the benefits and costs of achieving a rhythmic code based on aperiodic sensory measurements. We assume in this analysis a rate code, and fast synaptic transmission.

We presume there *is* a cost because moving from aperiodic sensory signals to rhythmic rate codes is a translation, and translations are rarely free from error. Several lines of evidence suggest oscillations in the central nervous system are non-stationary, what we will call bursty, (1–4), where band-power fluctuates on sub-second intervals. We suppose these bursts are functional or adaptive in some sense.

The benefits of a stationary oscillatory code have been frequently remarked on (5–27). Decades of empirical and theoretical work make it clear oscillations can modulate excitability (28), attention (27, 29) and create synchronous windows of neuronal communication (30). By temporally grouping action potentials, signal-to-noise increases (31), as does the number of coincident firing events (32, 33), which in turn drives learning at individual synapses (34–36). Most important here is the well-established knowledge that oscillations can modulate population firing rates (8, 10, 37). Here we argue any benefits must be weighed against any translation costs that come from adopting a rhythm code in the first place.

In this paper, we consider two modes of translation. First we study a Linear-Nonlinear-Poisson model (LNP) to model the effect of an global oscillator on independent units (38) (Figure 1a). Second, we examine recurrent biophysical circuits tuned to generate gamma oscillations that mimic human recordings (33) (Figure 1b). For both modes we measured translation costs and benefits using information theory. That is, we view oscillations as a communication channel.

**Fig. 1.**
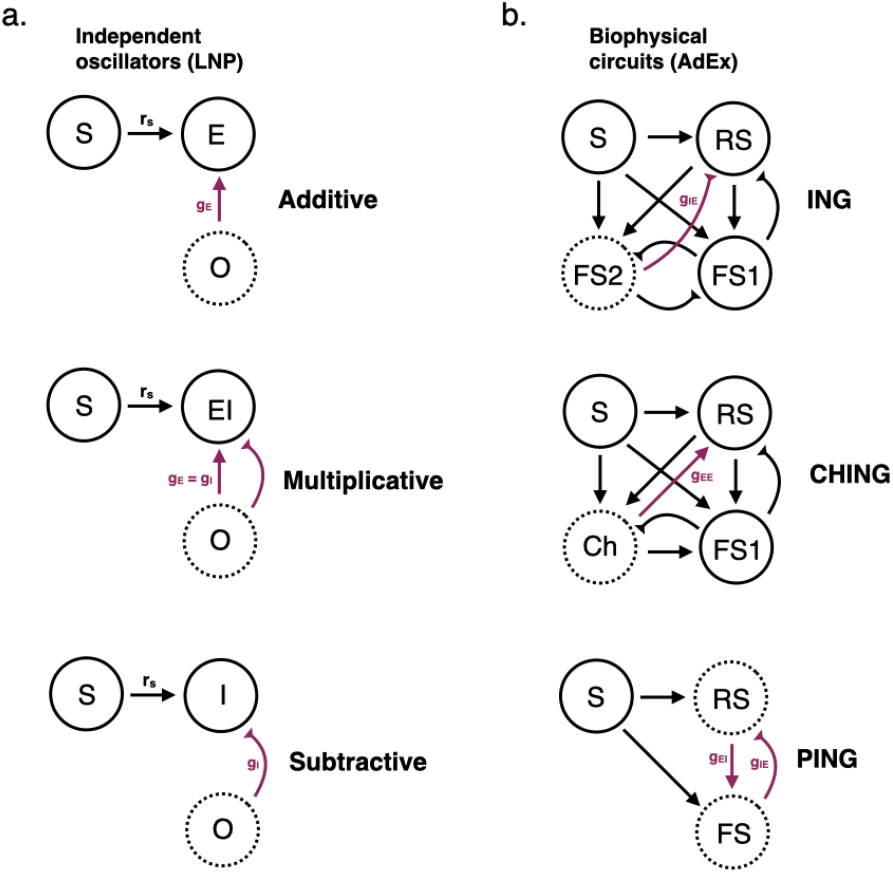
Network diagrams. **a**. Independent oscillators – *E, EI* and *I. Note:* Model II is minor variation in Model *I*, and is not shown. **b**. Biophysical circuits – ING, PING and CHING. Note that all connections are shown but only those labelled in purple are used to control rhythmic entrainment (directly) in our experiments here. Arrow points represent excitatory connections, while inhibitory connections are denoted by blunted heads. Dotted lines denote the oscillatory drive. Note how in panel a. The oscillator is uncoupled from the stimulus, and has no feedback with target populations (E,EI,I). In *b*. coupling between stimulus and all populations is complete.

We are surprised to report that no one physiological factor predicted the costs, or benefits, we observed by translating from an aperiodic to a rhythmic rate code. The circuit details, or translation mechanism, is a critical factor. So is the stimulus drive. Likewise the population size to-be-decoded has as strong an effect as the internal mechanisms themselves. To predict a burst as costly we must know a great deal about both its external drive, internal connections, and size of the population who will detect its message. These simulations suggest frequency and power are not enough to predict costs or benefits. We need to understand exacting physical details.

## Results

We simulated rate coding and oscillatory translation. First we used a LNP system (Linear-Nonlinear Poisson) of theta oscillations. This modeled global entrainment without feedback between the oscillation and the translated target. Second, we used a validated biophysical circuit model of human gamma rhythms. This second approach allowed for recurrent feedback between oscillators(s) and the target.

### Linear-Nonlinear Poisson and theta oscillations

Despite the oversimplifications present in all LNP models, they have proven remarkably useful in developing new theoretical ideas (39, 40). In our LNP model, a time-varying aperiodic stimulus is modulated by a theta oscillation. The result is nonlinearly filtered and Poisson sampled by independent units without intrinsic connections. We studied four kinds of modulation each made using the same global oscillator and independent population of neurons.

1. Model E – additive modulation, mimicking direct excitatory entrainment by AMPAergic synapses
2. Model I – subtractive modulation, mimicking direct inhibitory entrainment by GABAergic synapses
3. Model EI – multiplicative modulation, mimicking the gain effect of balanced microcircuits (41).
4. Model II – mimicked dual dendritic effects of inhibition, which can be both subtractive and divisive at the same time (42, 43).

Our goal in these first simulations was ape an oscillator modulating a rate code, where the oscillator and a single target population don’t share (immediate) recurrent connections (28, 37).

We measured conversion cost/benefit by making two information theory calculations. We first measured the information shared between a time-varying reference stimulus *s* (whose creation is described in the *Methods*) and a Poisson sample of that stimulus *S*. This quantifies the innate cost of translating a stimulus to a rate code by itself alone. In other words, this is a control condition. This control is the basis for our information measurements in Eq. 1, which we will now explain.

Let *S* be the Poisson rate time course for the control measure. And let it be given by 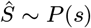 where *s* is the original target, and *P*(.) is a Poisson distribution. We use *r_S_*, etc, to denote the firing rate of samples from *P*. This sampling to give *S* then gives us our first mutual information value, MI(*s, S*). For the second we generated a “translated” time course *M*, using one of the models above (Figure 1**a**). With *M* we can measure the mutual information between *s* and *M*, giving MI(*s, M*), and our final equation.

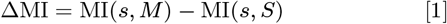

Examples of *s, S* and all models *M* are shown in Figure 2. The conversion from time courses to distribution, and a visual depiction of overlap between S and M is shown in Fig 3 and Fig 4, respectability.

**Fig. 2.**
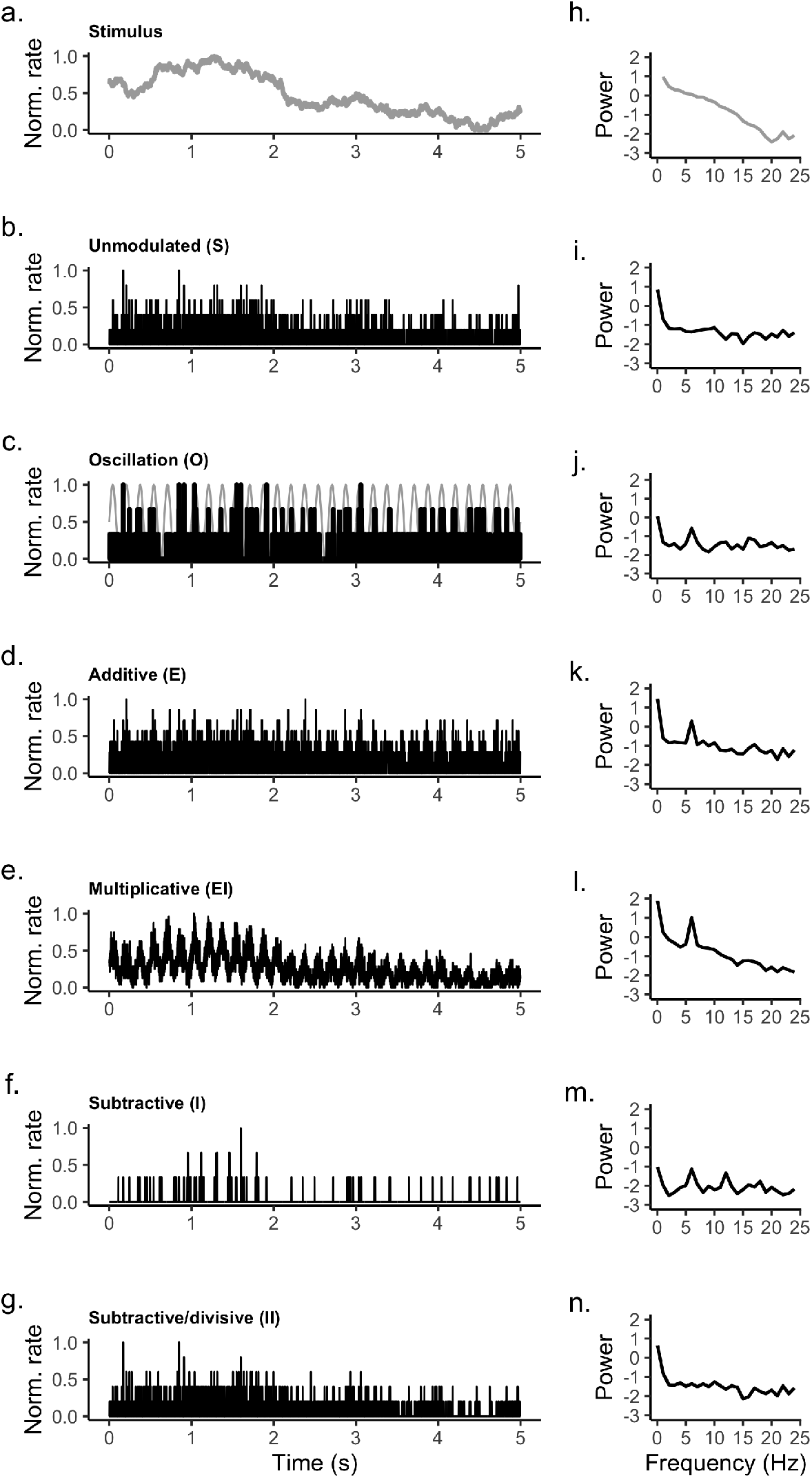
Examples of theta rhythm transformations in an LNP model. The left panels **f**-**j** are power spectra derived from the spiking time courses on the left. **a.** Example of a typical driving stimulus. **b.** A Poisson sampling of the (unmodulated) control stimulus, which serves as the non-oscillatory control condition for studying encoding between the stimulus and neural response, under a rate-coding regime **c.** The oscillation (grey) and a Poisson sampling of the same (black). This is denoted as model **O**. The remaining panels (d-g) show examples of the different kinds of transformation we studied here (38), where the Poisson rate coding system is modulated by different forms of oscillation. Note that all series were normalized between 0-1 in this figure to simplify the comparisons. *Simulation parameters:* the average stimulus firing rate *r_s_* was set to 10 spks/s, the oscillation average rate *r_o_* was 2 spks/s, *g* was 2, and the population size was 40. The oscillation’s oscillation frequency was 6 Hz.

**Fig. 3.**
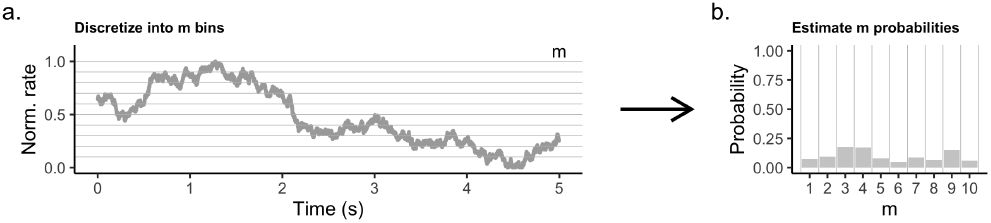
Step 1: estimating mutual information. We used a binning procedure to estimate discrete probability distributions, with *m* bins. **a**. Example of binning, using reference stimulus from Figure 3. **b**. The probability distribution derived from panel *a*. *Note*: The bin for the reference stimulus are represented as grey lines, and the model result is in black. Though LNP results are shown here, mutual information estimate and analysis were identical in both LNP and circuit models (shown below).

**Fig. 4.**
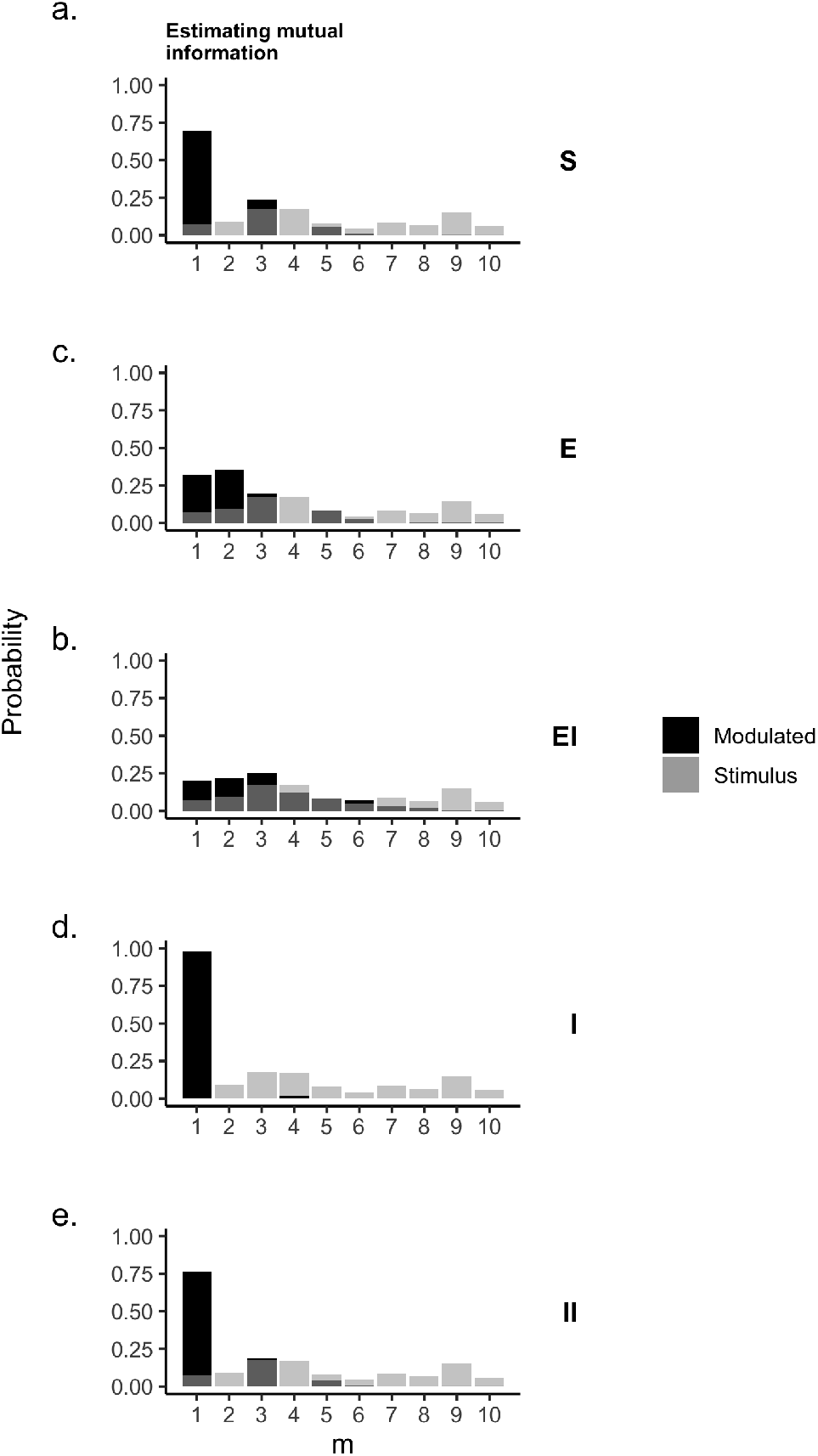
Step 2: estimating mutual information. Light grey is the stimulus distribution. Dark grey is the Poisson sample distribution. Overlap between the two light and dark provides a visual estimate of their mutual information (MI). That is, the more overlap, the higher their MI. **a.** An example distribution of the control model, *S*. **b-d**. Example distributions for all the transform/modulations. These distributions correspond to the timecourses shown in Figure 2. *Note*: Though LNP results are shown here, mutual information estimates and analysis were identical in both LNP and circuit models.

If Eq. 1 gives a negative value, this implies a translation cost. If it gives a positive value, this implies a translation benefit.

We’ll now review the translation benefits and costs we observed. The variables of foremost interest were the stimulus firing rate, the oscillation firing rate, the synaptic strength *g*, and the size of the population used to measure information. Other variables which proved less important are discussed later on.

The simple system we study first requires significant caveats to interpret. We’ll return to these in the discussion. Despite the limits imposed by such (over-)simplicity, the cost/benefit of translations here are nuanced.

We focus first on the effect of population size. If a few cells transmit a translated message this might be very different then if many cells do. Our interest in population size is based on the idea that even if a large number of neurons are entrained by an oscillation, many of those cells will in turn send messages to different target areas downstream. Cell numbers in these projections can be quite variable (44–47). So when we study population size, what we aim to measure is the effect of different “functional” population sizes on downstream targets.

“Small” populations of fewer than approximately 100 cells lead to better outcomes, overall for all LNP models. For excitatory modulations (E/EI) translation benefit would be predicted to peak below 100 cells (Figure 5**b**). For inhibitory modulations (I/II) the predicted translation cost is minimized below 100 cells (Figure 5**b**. Owing to the importance of population size, we will focus on a “small” population of 40 cells, and “large” population of 10240 cells in this section.

**Fig. 5.**
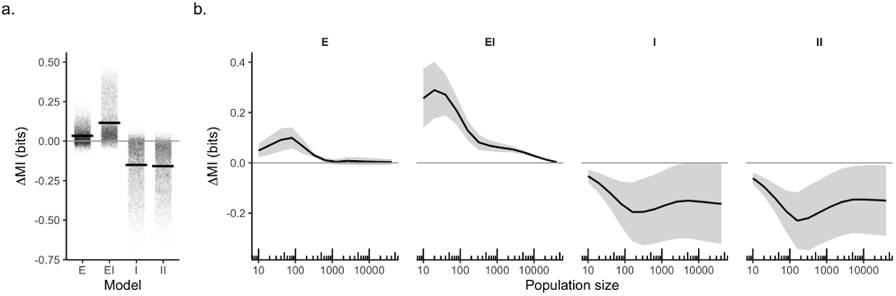
The importance of population size, with an independent oscillator. **a)** Total change in mutual information (ΔMI), across all population sizes, for all four LNP models. Individual points represent individual trials (*n*=100). Black bar is the average. **e)** The average change in mutual information (ΔMI) plotted as a function of population size. Error bars represent standard deviation. *Simulation parameters:* the average stimulus firing rate *r_s_* was set to 10 spks/s, the average oscillation rate *r_o_* was 2 spks/s, *g* ranged from 1-8 in one unit increments. The oscillation frequency was 6 Hz.

In the small population Model E (additive translation), we report an average benefit of about 0.1 bits over the control (Figure 8**a**). This decreases with stimulus firing rate (Figure 8**b**) and increases with both the oscillation firing rate (Figure 8**b**) and the synaptic strength *g* (Figure **8,** all panels).

**Fig. 6.**
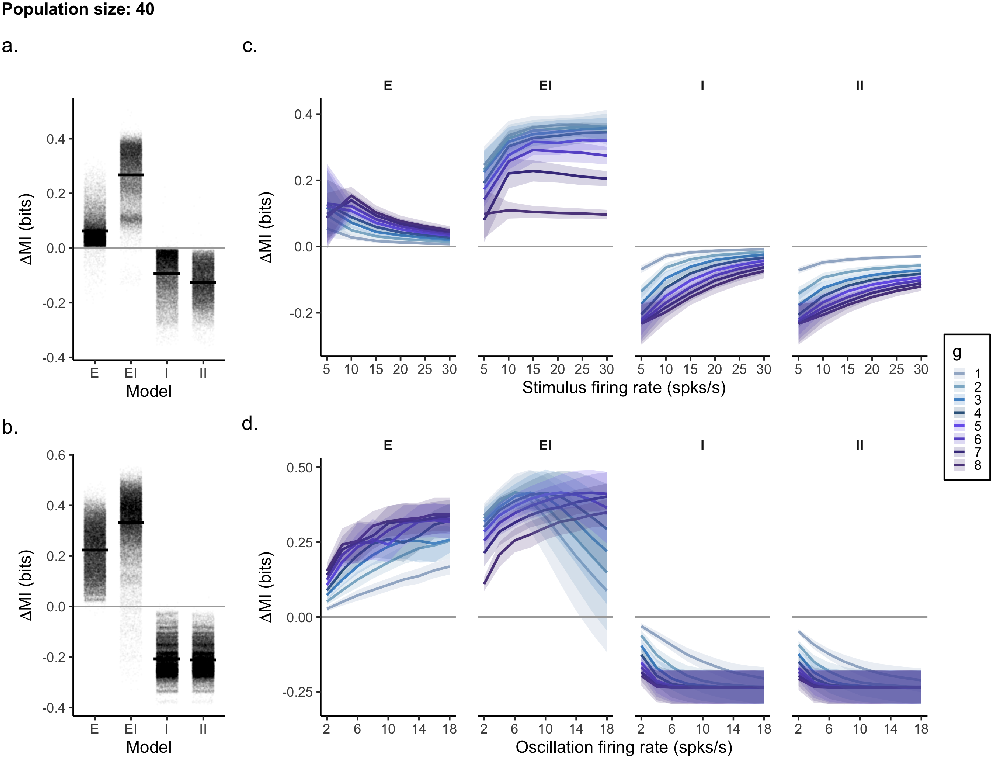
The effect of firing rate on *small* population sizes, with an independent oscillator. Top panels explore stimulus firing rate. Bottom panels explore the effect of the oscillation’s firing rate. **a),c** Total change in mutual information (ΔMI), across all conditions as a function of firing rate. Individual points represent individual trials. Black bar is the average. **b),d** The average change in mutual information with oscillation’s firing rate and relative synaptic strength *g* (blue). Error bars represent standard deviation. *Simulation parameters*: the average stimulus firing rate *r_s_* ranged from 5-30 spks/s, the oscillation’s average firing rate *r_o_* ranged from 2-18 spks/s, *g* ranged from 1-8. The oscillation frequency was 6 Hz.

**Fig. 7.**
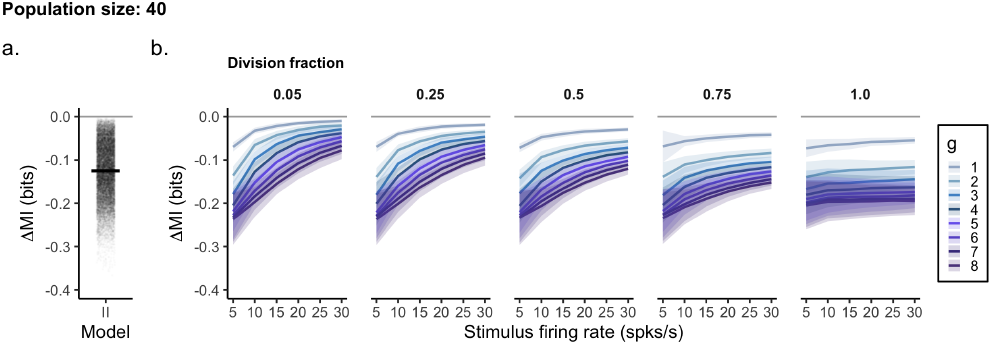
Stimulus firing rate and its effect on oscillatory encoding in Model **II**. **a)** Total change in mutual information (ΔMI). Individual points represent individual trials. Black bar is the average. **e)** Average change in mutual information (ΔMI) plotted as a function of stimulus firing firing rate, and relative synaptic strength *g* (blue) *Simulation parameters:* the average stimulus firing rate *r_s_* ranged from 5-30 spks/s, the oscillation’s average firing rate *r_o_* was 2 spks/s, *g* ranged from 1-8. The oscillation frequency was 6 Hz.

**Fig. 8.**
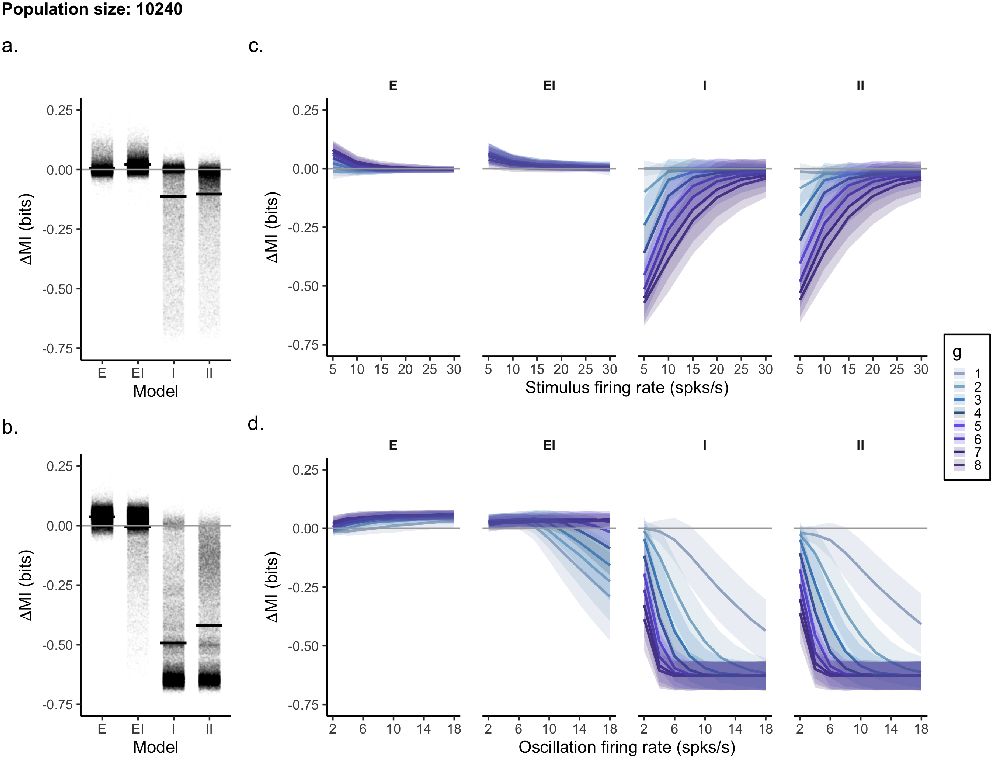
The effect of firing rate on *large* population sizes with an independent oscillator. Top panels explore stimulus firing rate. Bottom panels explore the effect of the oscillation’s firing rate. **a),c** Total change in mutual information (ΔMI), across all conditions. Individual points represent individual trials. **b),d** The average change in mutual information with firing rate and relative synaptic strength *g* (blue). Error bars represent standard deviation. *Simulation parameters:* the average stimulus firing rate *r_s_* ranged from 5-30 spks/s, the oscillation’s average firing rate *r_o_* ranged from 2-18 spks/s, *g* ranged from 1-8. The oscillation frequency was 6 Hz.

In the small population with Model EI (multiplicative translation) we report an average benefit of about 0.25 bits (Figure 8**a**). This scaled with both the stimulus and oscillation firing rates (Figure 8**b-d**), until the effect plateaued. As expected for any model of balanced modulation we saw an inverse relationship between synaptic strength and mutual information (41). For weak synapses and strong oscillatory firing we observed a decrease in benefit once the oscillation’s firing rate exceeded the stimulus rate. This is 10 spks/s in Figure 8.

With both kinds of inhibitory modulation *I* and *II* we report a translation cost, in most conditions. This had an average value of about −0.1 bits (Figure 5**a**; Figure 8). The cost was to some degree mitigated by the stimulus strength (Figure 8**b**), and made worse by the oscillation’s firing rate and synaptic strength. In model II, the ratio of divisive to subtractive inhibition did not affect the overall cost. Extensive divisive inhibition did however prevent strong stimulus rates from mitigating the translation cost (Figure 5**b**).

The general lesson from our theta/LNP simulation is this: there are working ranges where oscillations are beneficial for communication: when those oscillations can correct for “undersampling” of the stimulus. That is, they “add” spikes in a way that lets the population more closely resemble the stimulus. This phenomenon is deeply related to prior modeling of how deep brain stimulation can improve neural functioning (48). However, as soon as the population is large enough to sample adequately on its own, or if the oscillation’s influence grows to large then this benefit is overcome by the oscillation’s acting to instead distort the message.

#### Control experiments

##### Oscillation frequency

We have studied a theta 6 Hz oscillation so far in our LNP approach. When we examined frequency across a typical physiological range, we observed few changes to our results (Figure 9**a**). This comes from our choice of rate models, and linear translation. We would expect a very different result for a biophysical model, or one that considers spike-timing, rather than just rate coding.

**Fig. 9.**
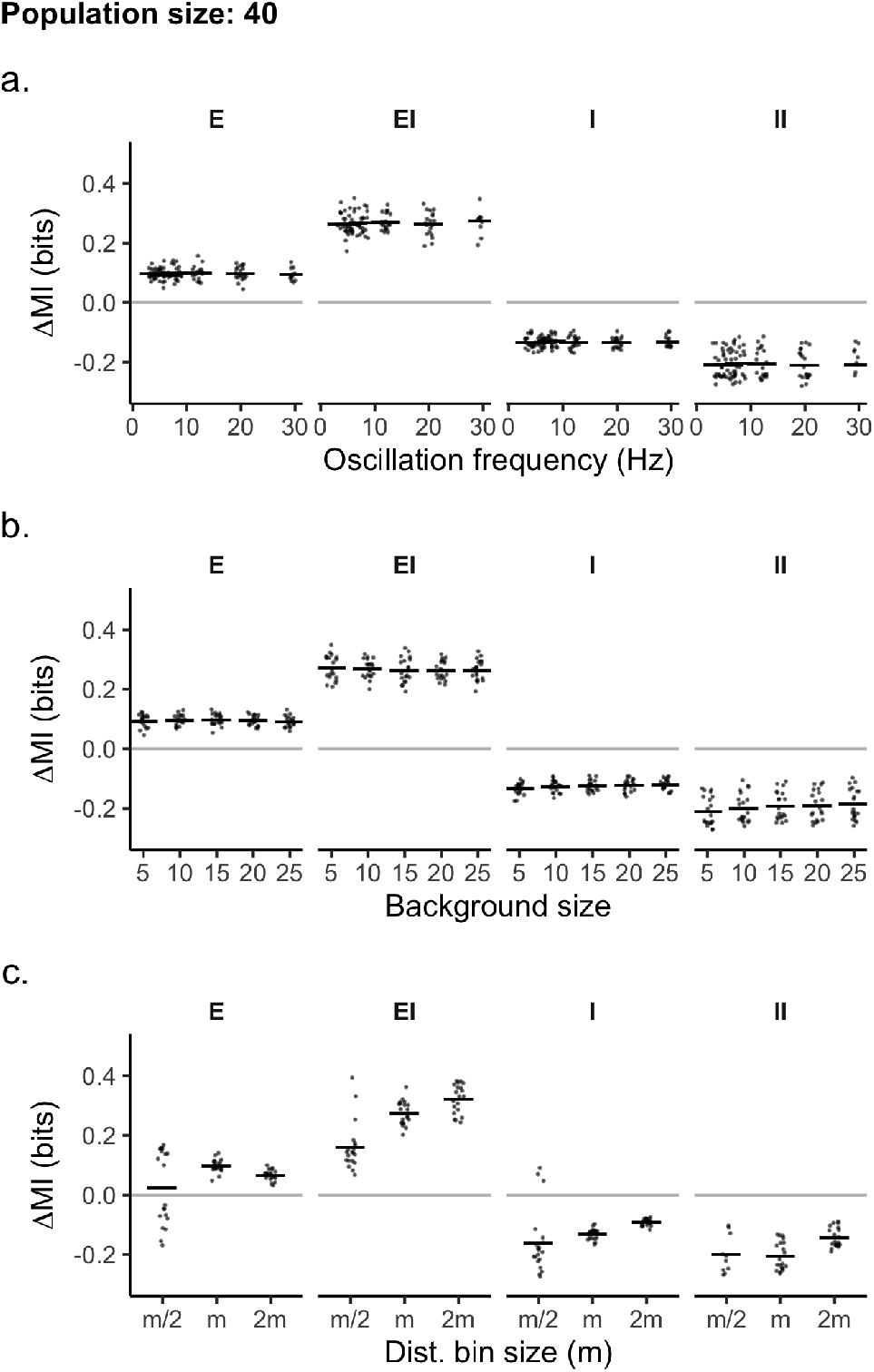
**a**. Oscillation frequency and its own effect on oscillatory encoding. **b**. Background population size and its effect on oscillatory encoding. **c**. Bin size and its effect on oscillatory encoding.

##### Background noise (size)

We have simulated a background noise level of about 2 spikes/s for all previous simulations, which is consistent with most cortical recordings Varying this more than 10 fold did not affect the benefits or costs we observed (Figure 9**c**).

##### Bin size

Our calculation of mutual information relies on binned data, and our first choice for the bin number (*m*) was to have it match the average stimulus firing rate. This was about 10 spikes/s. This makes our entropy calculations have a consistent 1 spks/s resolution. Doubling or halving our bin number had no qualitative effect on the results we observed (Figure 9**d**). This seems to validate our choice.

### Biophysical circuit

We studied a biophysical model of gamma oscillations, tuned to match human physiology (33). The sensory stimulus we used to drive activity was identical to our LNP mode. As was our general analysis method. We studied three kinds of gamma circuits, ING, PING, and CHING, each made using many of the same baseline parameters (Table 2-3). Examples of model output are shown in Figure 10.

**Fig. 10.**
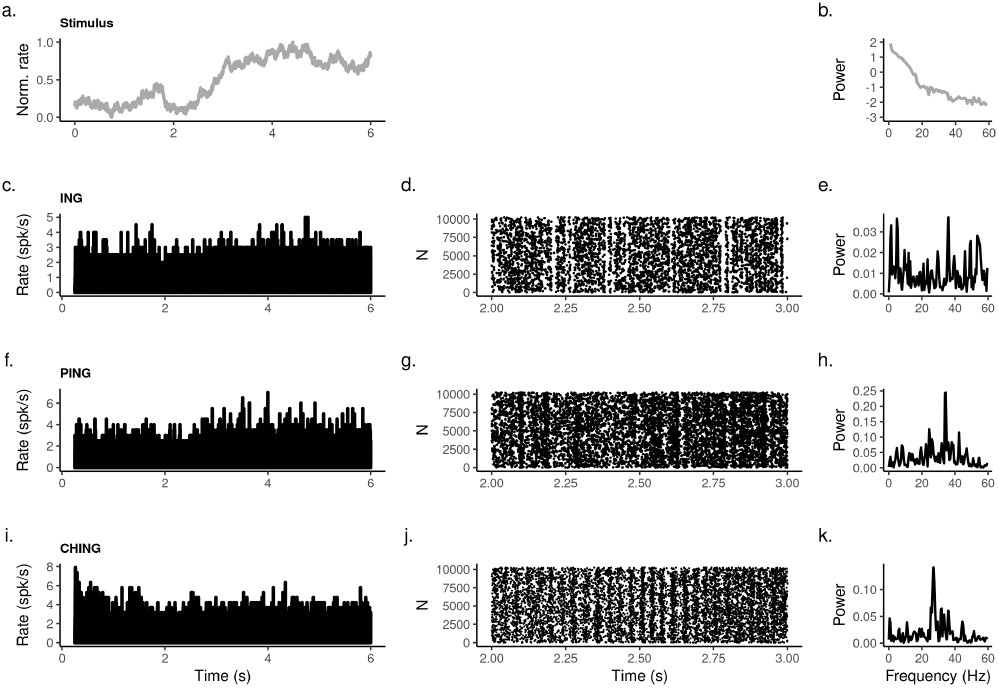
Examples of rhythmic transformations in a biophysical model. The left panels **f**-**j** are power spectra derived from the spiking time courses on the right. In the center column is a raster of spike-times over a 1 second window that illuminates the “fast” oscillation more clearly. **a.** Example of a typical driving stimulus. The remaining panels (*c-k*) show examples of the different kinds of transformations for our three network models of human gamma oscillations. *Simulation parameters:* the average stimulus firing rate *r_s_* was set to 1 spks/s, synaptic conductances and time constants were set to their default values (See Table 3. The population size was 10240.

**Table 1.** Model parameters, and general mathematical terms, for the independent oscillator. When appropriate translation parameter ranges are listed first, with a typical value shown in parenthesis shown in parenthesis after the range.

| Term | Description | Value or formula |
| --- | --- | --- |
| B | the background drive | - |
| $r_b$ | the average firing rate of B | 2 |
| S | the stimulus drive | - |
| $r_s$ | the average firing rate of $s$ | 2 – 30 (10) |
| $\sigma_r$ | variance of $s$ | 0.01$r_s$ |
| $\hat{S}$ | Poisson firing; a sample of S | Eq. 6 |
| O | the oscillating drive | - |
| $r_o$ | the average firing rate of O | 2 – 8 (2) |
| $f$ | oscillation frequency | 4 – 30 (6) |
| $\hat{O}$ | Poisson firing; a sample of O | Eq. 6 |
| $g$ | modulation strength | 1 – 8 |
| E | additive modulation (a drive) | Eq. 7 |
| I | subtractive modulation (a drive) | Eq. 8 |
| EI | multiplicative modulation (a drive) | Eq. 9 |
| II | subtractive/division modulation (a drive) | Eq. 10 |
| $\hat{E}$ | Poisson firing; a sample of E | Eq. 6 |
| $\hat{I}$ | Poisson firing; a sample of I | Eq. 6 |
| $\hat{EI}$ | Poisson firing; a sample of EI | Eq. 6 |
| $\hat{II}$ | Poisson firing; a sample of II | Eq. 6 |
| $m$ | Bin size | $r_s$ |
| H | Shannon entropy | Eq. 17 |
| MI | mutual information | Eq. 18 |
| $N(0, 1)$ | a normal sample | - |
| $\tau_\alpha$ | the alpha kernel | TODO |

**Table 2.**
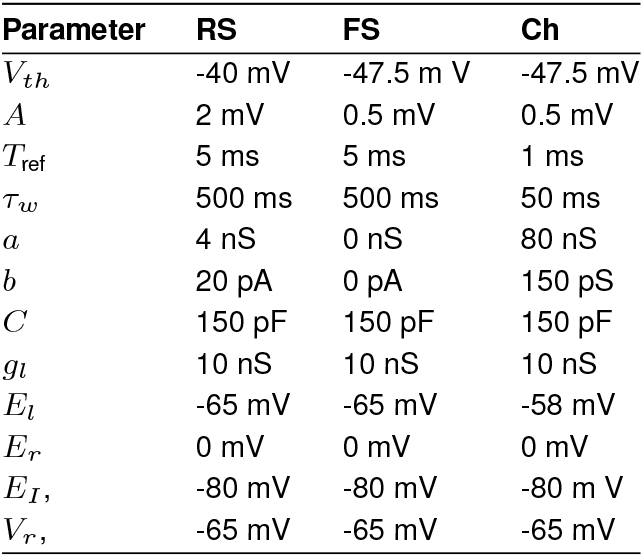
Population parameters for the biophysical network, with populations of fasting spiking cells (FS), regular spiking cells (FS) and chattering cells (Ch).

**Table 3.**
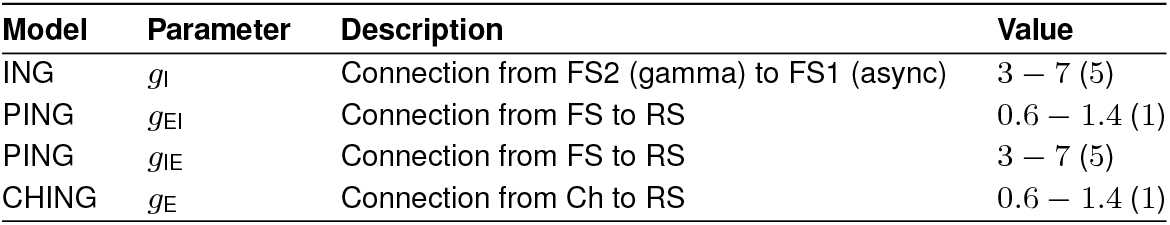
Translation parameters for populations in the biophysical circuit. Parameter ranges are listed first with the typical value, when appropriate, shown in parenthesis after the range. Values are in nS.

In each of these models there is a dependence between the oscillatory generator, the driving stimulus, and the whole recurrent circuits. These models are based on those of Susin and Destexhe (49), and tuned to match human 20-80 Hz (gamma) oscillations. This is the sort of recurrent connections found in cortical microcircuits (47, 50, 51), though simplified.

1. Model ING – modulation of a coupled E-I population by an inhibitory gamma rhythm, mimicking recurrent inhibitory entrainment by GABAergic synapses.
2. Model CHING – modulation of a coupled E-I population by an excitatory gamma rhythm, mimicking recurrent excitatory entrainment by AMPAergic synapses
3. Model PING – a coupled E-I population that self-generated a gamma rhythm.

We’ll now review the translation benefits and costs we observed. The variables of foremost interest were the stimulus firing rate, select synaptic strengths *g*, and the size of the population. Not unlike the first model, units in the conductance terms for each circuit are critical values. Synaptic time constants were generally not critical, with one exception.

In two of the three circuits we studied “small” populations of between 100-1000 cells lead to better outcomes, overall. As did comparatively weak stimulation in all circuits. However the other, ING, and inhibitory oscillation, showed an increase in information flow with population size which is the opposite pattern seen in the independent oscillator.

Owing to the importance of population size, we will still focus for the remainder of this paper on a “small” populations of either todo or todo cells, and a “larger” population of 10240 cells.

In the small population in Model ING (inhibitory-cell gamma) we report an average cost of about −0.05 bits over the control (Figure 11**a**). For weak stimuli observed a benefit of about 0.25, depending on *g*. This decreases with stimulus firing rate (Figure 11**b**, top left panel), but increases with both the synaptic strength *g*.

**Fig. 11.**
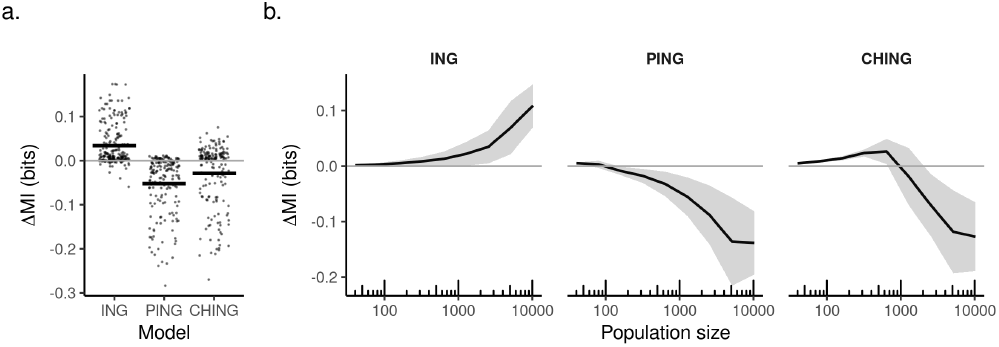
The (continuing) importance of population size when decoding a recurrent circuit. **a)** Total change in mutual information (ΔMI), across all population sizes, for all three circuit models. Individual points represent individual trials (*n*=20). Black bar is the average. **e)** The average change in mutual information (ΔMI) plotted as a function of population size. Error bars represent standard deviation. *Simulation parameters:* the average stimulus firing rate *r_s_* was set to 1 spks/s, the different *g* for each circuit were set to ….

In the small population with Model CHING (chattering-cell excitatory gamma) we report a net of cost of −0.25 bits (Figure 5**a**). This scaled negatively with both the stimulus (Figure 5**b**, top right panel), until the effect plateaued. We saw an inverse relationship between synaptic strength and mutual information. For weak synapses and weak stimuli we observed an average benefit of about 0.1 bits (Figure 5**b**, top right panel).

When we searched for conductance and stimulus interactions in Model PING (*EtoI* and *ItoE*) we reported a net translation cost of about −0.3 bits (Figure 5**a**). The only exception is a weak (0.5 spks/s) stimulus, combined with large relative conductance values (1.4 or 7 nS) (Figure 5**b**, bottom panels).

For both population size, and across all three model types, there was little dependence on information flow and the choice of synaptic time constant. The exception to this rule was in Model PING and the excitatory constant. Here costly oscillations could be converted to beneficial ones as the time constant rose. This effect was most strong for the “small” population. The origin of this particular effect is unclear. What is clear is that unlike the case of oscillatory resonance, time constants are not central to understanding oscillatory translations. At least, in this circuit.

The general lesson from our beta/gamma simulation is this: only inhibitory entrainment, and its contaminant generation of network-level spiking events (18, 52) can benefit information transmission as have defined it here. That is, they “add” spikes in way that lets the population more closely resemble the stimulus.

### Average, best, and worst case analysis

We have characterized rhythmic encoding by assuming the average case is the most relevant. And from the point of view of statistical inference that is often so. But from the point of view of computer science it is often fruitful to consider two other cases: the worst case outcome and the best case outcome. This is because sometimes errors can come with substantial outside consequences. In such cases a worst case disposition is appropriate. While in other situations errors can be inconsequential, and so the focus is on the best case.

For many information flow studies the default choice has been the average case. We argue the disposition one takes should depend on the beliefs one has about the uniqueness or degeneracy of the message. In other words, the case analysis should depend on how central a (translated) population is to the larger distributed computation. Concrete examples of this kind might be oscillations and information flowing from either thalamus, or hippocampus. Both regions play central roles in routing sensory information (37) and controlling memory (53, 54).

If a population being translated is providing unique information, then worst case analysis should apply. In this case translation errors might translate directly to behavioral errors. If however the population being translated is, in effect, a degenerate or partial “voice”, then best-case analysis may be the right choice. Here any translation errors which are introduced could be corrected. While any improvements would serve to bootstrap up the whole computation? Examples of this might be cortical-cortical oscillations, as broadly speaking information storage in the cortex seems to itself both sparse and degenerate (55, 56).

The average case can be seen perhaps as a safe middle ground. When we do not have beliefs about how unique or degenerate a “message” might be, it is best to choose the unbiased average. In other words, the standard approach may be the safest but also, as we now show, safe can be rather misleading.

When we consider these three different “dispositions”–average, best, and worst–the results we can report change radically.

In Figure **15a-b** we re-plot the average case from the LNP model in Figure 5 **b**. We then compare this to both the min (Figure **15c-d**) and max (Figure **15e-f**) taken from the same experiment. In the worst case oscillations are a net cost across all conditions. In the best case they are a net gain across all conditions. However these best and worst cases are not uniform across oscillation physiological conditions.

**Fig. 12.**
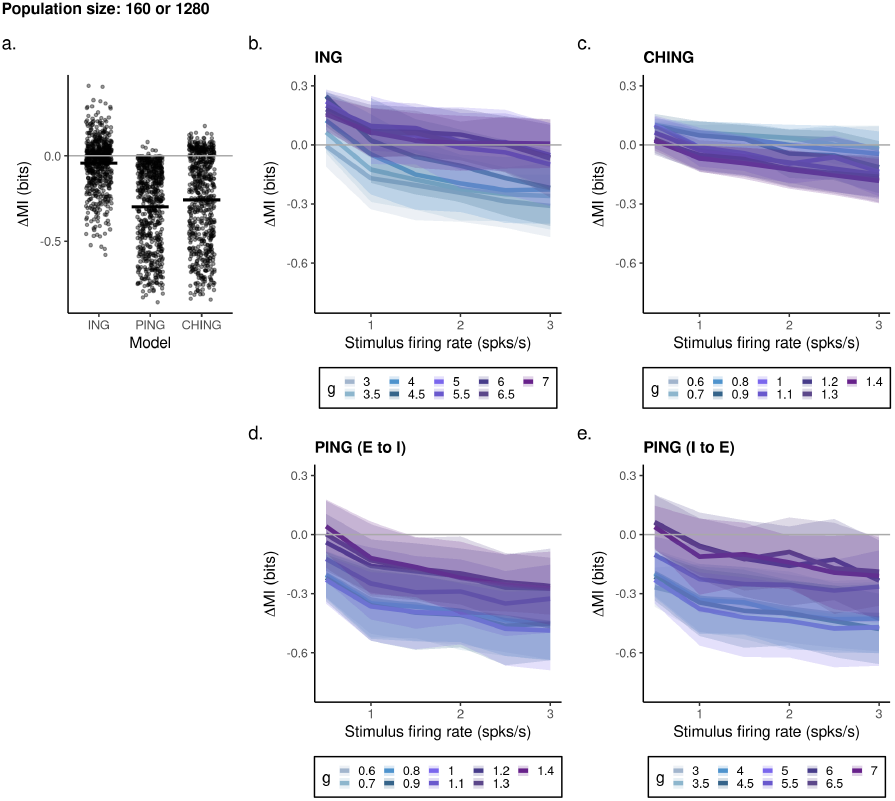
The effect of stimulus firing and synaptic conductance on *small* population sizes in three recurrent circuits. **a** Total change in mutual information (ΔMI), across all conditions as a function of all firing rates. Individual points represent individual trials. Black bar is the average. **b),d** The average change in mutual information with changes in stimulus firing rate and relative synaptic strength *g* (blue). Error bars represent standard deviation. *Simulation parameters:* the average stimulus firing rate *r_s_* ranged from 0.5-3.0 spks/s, *g* ranged from 0.6-7.0 nS (see Figure legends for circuit specific values).

**Fig. 13.**
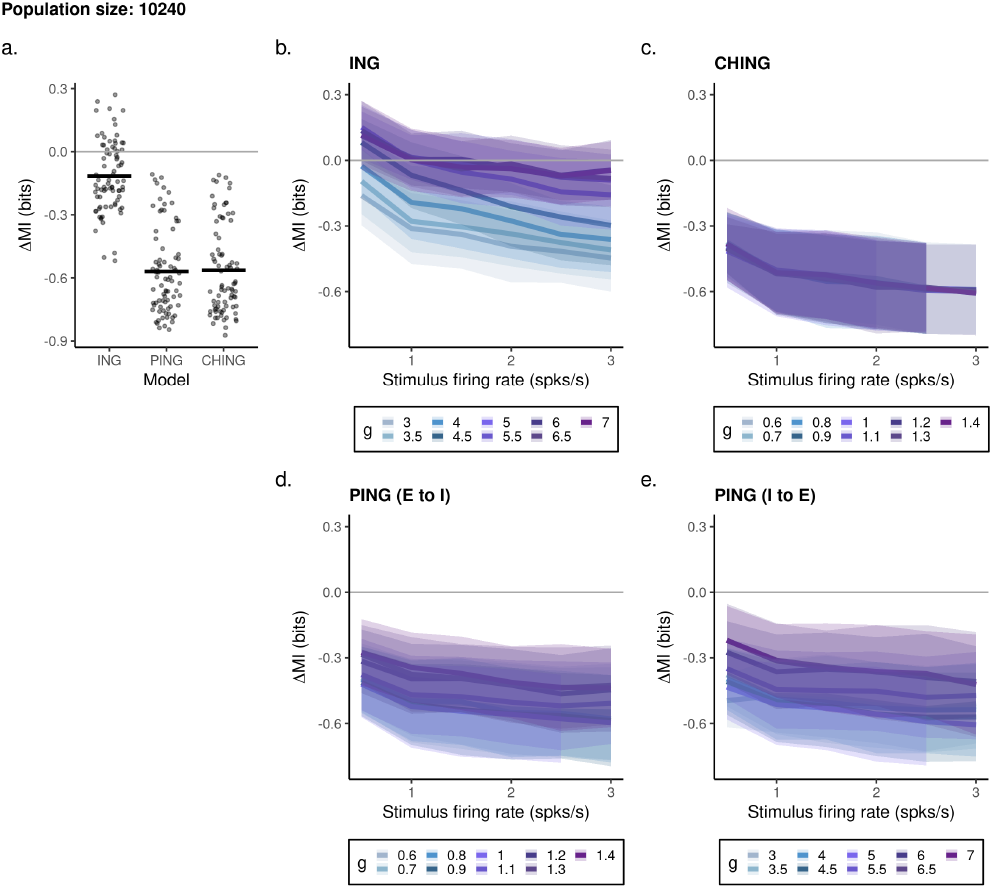
The effect of stimulus firing synaptic conductance on *large* population sizes in three recurrent circuits. **a** Total change in mutual information (ΔMI), across all conditions as a function of all firing rates. Individual points represent individual trials. Black bar is the average. **b),d** The average change in mutual information with changes in stimulus firing rate and relative synaptic strength *g* (blue). Error bars represent standard deviation. *Simulation parameters:* the average stimulus firing rate *r_s_* ranged from 0.5-3.0 spks/s, *g* ranged from 0.6-7.0 nS (see Figure legends for circuit specific values).

**Fig. 14.**
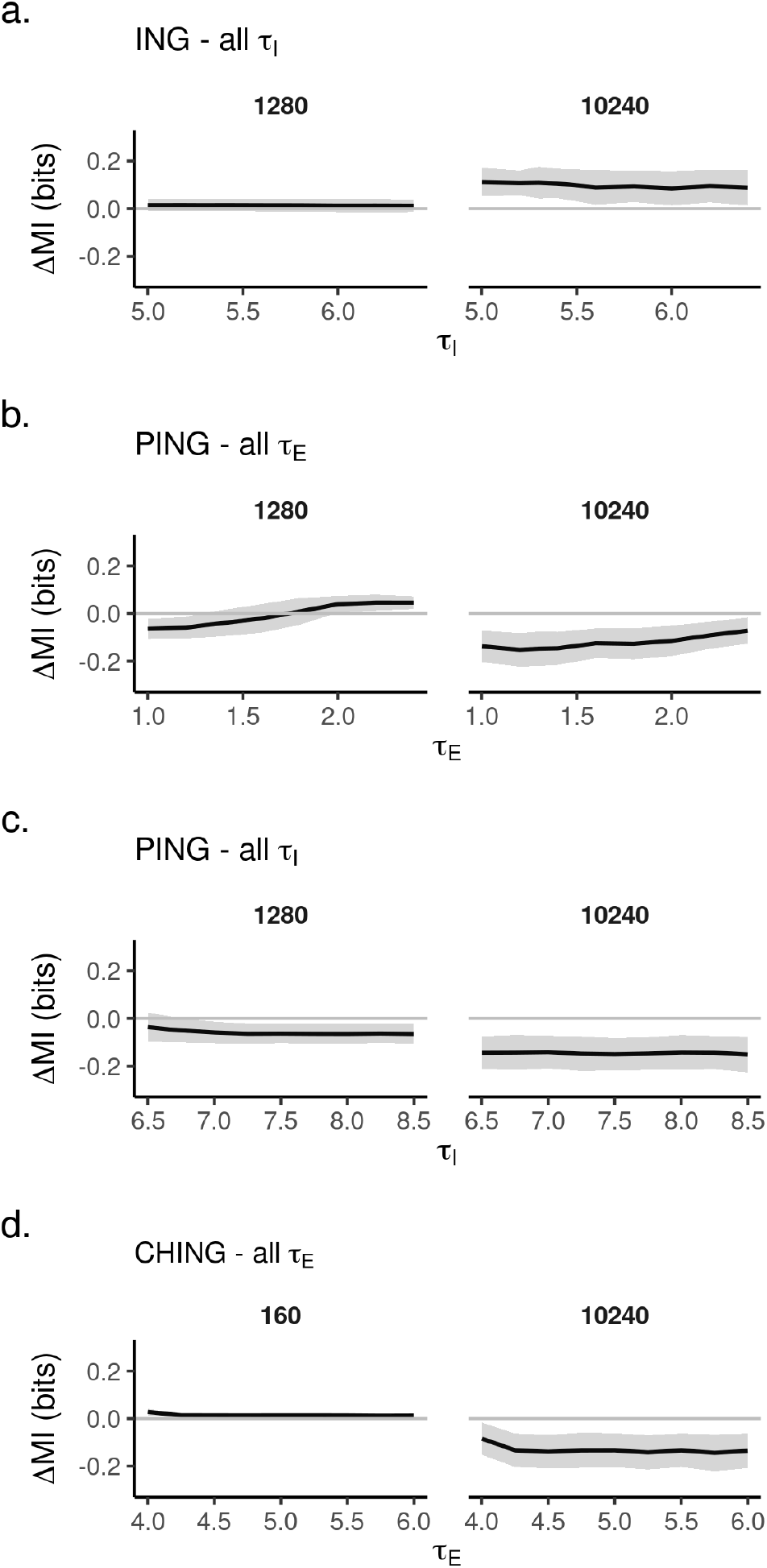
The effect of “fast” synaptic time-constants on in three recurrent circuits. All rows depict the average change in mutual information (ΔMI) with changes to synaptic time constants. Error bars represent standard deviation. Left column is the “small” population. Right is the “large” population. **a**. The effect of varying inhibitory constants on Model ING. **b**. The effect of varying excitatory constants on Model PING. **c**. The effect of varying inhibitory constants on Model PING. **e**. The effect of varying excitatory constants on Model CHING. *Simulation parameters:* the average stimulus firing rate *r_s_* was fixed at 1 spks/s, *τ* values are as shown, and all other parameters were set to default values (Table 2-3).

**Fig. 15.**
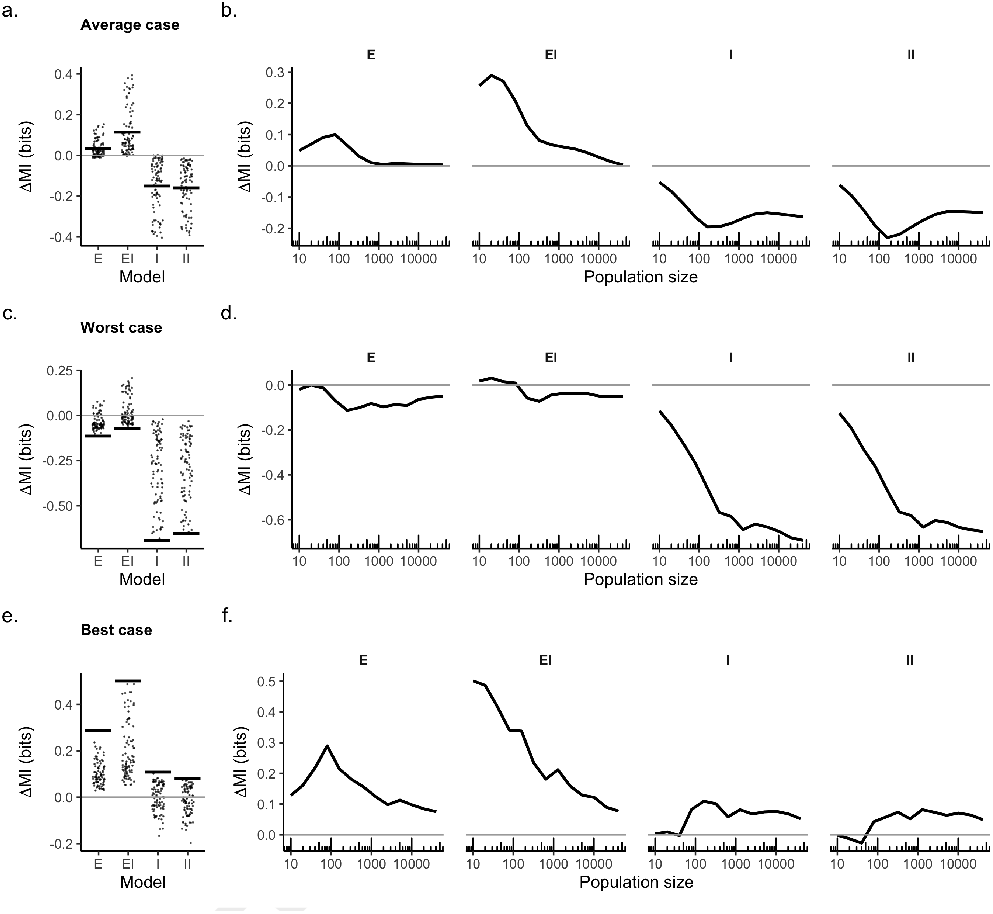
Average, best, and worst case analysis with the independent oscillator. **a-b)** Total change in mutual information in the *average case*. a.For each model, summed over population size and synaptic strength. *b*. As a function of population size. **c-d)** Total change in mutual information in the *worst case. c*. For each model, summed over population size and synaptic strength. *d*. As a function of population size. **e-f)** Total change in mutual information in the *best case. e*. For each model, summed over population size and synaptic strength. *f*. As a function of population size. *Simulation parameters:* the average stimulus firing rate *r_s_* was set to 10 spks/s, the oscillation’s average firing rate *r_o_* was 2 spks/s, *g* ranged from 1-8. The oscillation frequency was 6 Hz.

In Figure 16 re-examine our three circuit models. Here worst-case analysis continues to turn benefit conditions into costly ones, and make already costly translations worse. Bestcases continue to “rescue” costly average information cases.

**Fig. 16.**
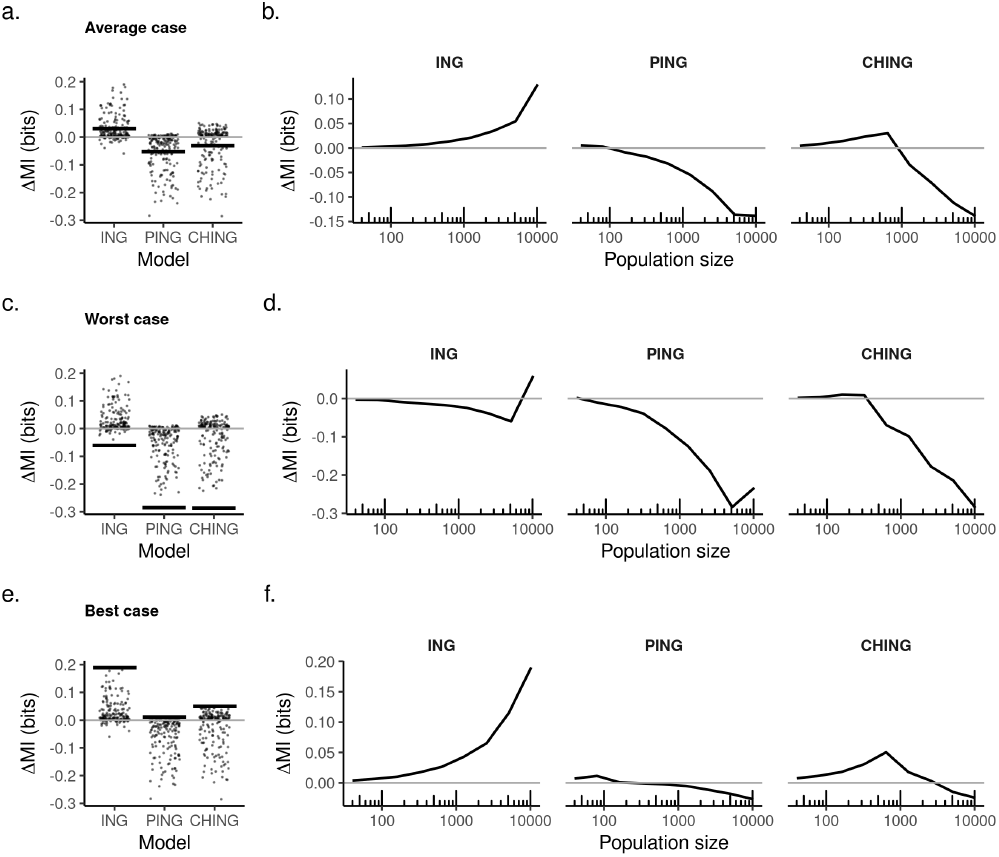
Average, best, and worst case analysis with the independent oscillator.. **a-b)** Total change in mutual information in the *average case*. *a*.For each model, summed over population size. *b*. As a function of population size. **c-d)** Total change in mutual information in the *worst case. c*. For each model, summed over population size. *d*. As a function of population size. **e-f)** Total change in mutual information in the *best case. e*. For each model, summed over population size. *f*. As a function of population size. *Simulation parameters:* the average stimulus firing rate *r_s_* was set to 1 spks/s, and all other parameters were set to default values (Table 2-3).

We are aware of no scientific work looking at the error “disposition” of neural communications. Given how severe it changes our results, and given how errors can matter little in play or substantially during precision tasks like surgery, we argue this kind of analysis is relevant, and under-studied. We hypothesize answers can be found by considering the potential uniqueness of the communication.

### Alternate error analysis

The use of distributions for time-domain data also destroys the essential temporal nature of the signals. This might be important here. We therefore re-ran select experiments from the biophysical circuit, but this time tabulated costs/benefits using euclidean errors calculated on a moment-by-moment basis (Eqs. 2-3). In Eq. 2, *T* is the total simulation time, *y* is some observed signal, and 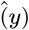 is some given target.

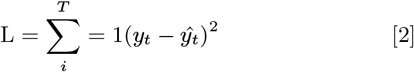

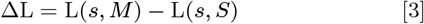

We report the same pattern of results using time-domain errors as we saw using information theory in the circuit model. Compare Figure 11 to Figure 17. If anything, the different patterns are more pronounced in information analysis, justifying our choice in using it.

**Fig. 17.**
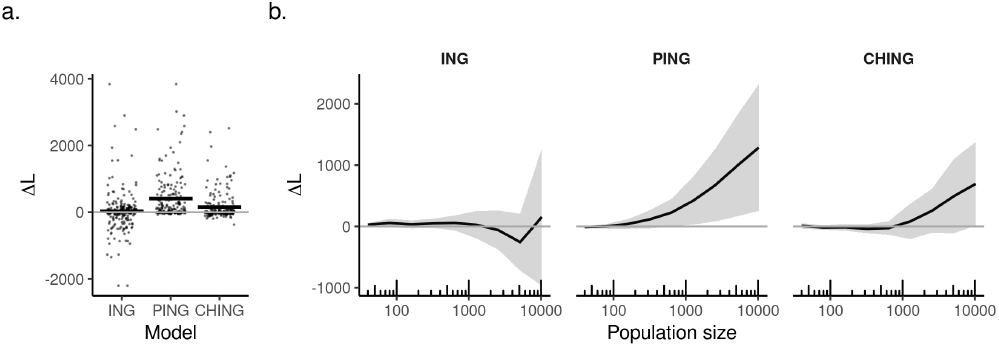
Alternative error analysis of population size when decoding a recurrent circuit. **a)** Total change in error (ΔL) across population sizes, for our three circuit models. Individual points represent individual trials (*n*=20). Black bar is the average. **e)** The average change in error (ΔL) plotted as a function of population size. Error bars represent standard deviation. *Simulation parameters*: the average stimulus firing rate *r_s_* was set to 1 spks/s. All other simulation values were set to default values (Table 2-3).

## Methods

The methods for this work are divided into parts. The simulations, and the information theory calculations.

### Simulations

We ran two kinds of simulations. One model featured an independent oscillator, and assumed a LNP model of the neural code. The second model was biophysical consisting of several neural populations in one of three recurrent arrangements.

#### Linear-Nonlinear Poisson

Each LNP model was composed of four parts. The driving stimulus, the oscillation, the target firing population, and the background noise.

To generate the stimulus we simulated a one dimensional diffusion process, as those have been suggested to act as a reasonable approximation to the “naturalistic” firing patterns observed in early visual areas during passive movie viewing (57). This is given by Equation 4, where *N*(0, 1) is a *sample* from the normal distribution which gets rescaled by *σ_s_* = 0.01*r_s_*, where *r_s_* is the average drive. The stimulus is further constrained such that 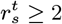. Note: when we refer to a value that is a function of time (*i.e., r_s_*(*t*)) we use the shorthand, 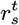. We set 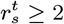 to avoid pauses during stimulation which can artificially inflate correlations/mutual information. If 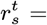 this will in some model settings lead to a silencing of activity of the population(s) as whole. This in turn induces a “false” correlation between the driver and the population.

In general we use use lower case *r* to refer to individual samples, *r* for average values, and uppercase letters, *S, M, E*, etc to refer or time course of values or to refer to the model classes themselves; their being synonymous for our purposes

Oscillatory firing was sinusoidal, and given by Equation 5. Where *f* is the periodicity in Hz. *r_o_* was fixed at 2 Hz, unless so noted. The third drive is, as noted, the background which is a constant 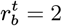.

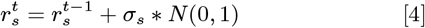

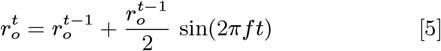

The oscillation rate 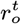 was modulated (or not) and results were independently sampled by *N* independent Poisson neurons. Prior to Poisson sampling, all firing drives were nonlinearly transformed by a threshold-linear activation function (38). The combination of linear interactions, nonlinearly filtering, and Poisson sampling is what marks our model as an LNP system (40).

Firing in the target population was based on a Poisson process. This is given by Equation 6, where *r* is the firing rate, or drive, and *k* is the number of spikes per time step (1 ms for all models here). Firing was limited to be no faster than once every 2 milliseconds, the absolute factory period.

For each trial in every experiment a new random seed was chosen. This generated a new naturalistic stimulus, as well as a new population of Poisson neurons. Every experiment consisted of 100 trials. Before analysis all spike trains for individual Poisson neurons were transformed to population average firing rate, denoted by a hat symbol. For example, 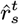 is the instantaneous Poisson firing rate from the stimulus drive 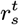. The temporal resolution used was 1 ms.

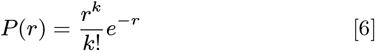

We assume that “synapses” for the stimulus into the target population were fixed, and unitary. This means the equations for modulation are as follows. The oscillation drive 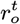 modulates the stimulus 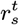 with a variable “synaptic strength” *g*. Modulation comes in one of four kinds – additive (Eq 7, subtractive (Eq 8, multiplicative (Eq 9, and subtractive/divisive (Eq 10. As a shorthand we refer to these as E, I, EI, and II respectively. Examples of each are shown in Figure 2.

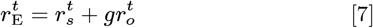

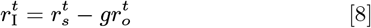

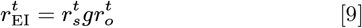

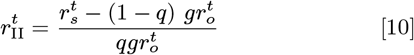

Our first model of inhibition **I** is perhaps too simple, given what is presently understood about the complexity of cortical inhibition (42, 58–60). This is because inhibition in the cortex, our prime area of interest, is targeted to both the dendrites and the soma. With recent work suggesting the soma experiences divisive inhibition at the soma, while dendrite targeting interneurons act subtractively. Papasavvas *et al* (43) developed a rate model of this two part inhibition scheme, and we adapted it to fit our LNP system. Here we implement a linear trade-off between the amount of divisive and subtractive inhibition, controlled by parameter *q* (Eq. 10).

Examples of all arithmetic codes are shown in Figure 18 where they are presented in terms of the classic FI-curve.

**Fig. 18.**
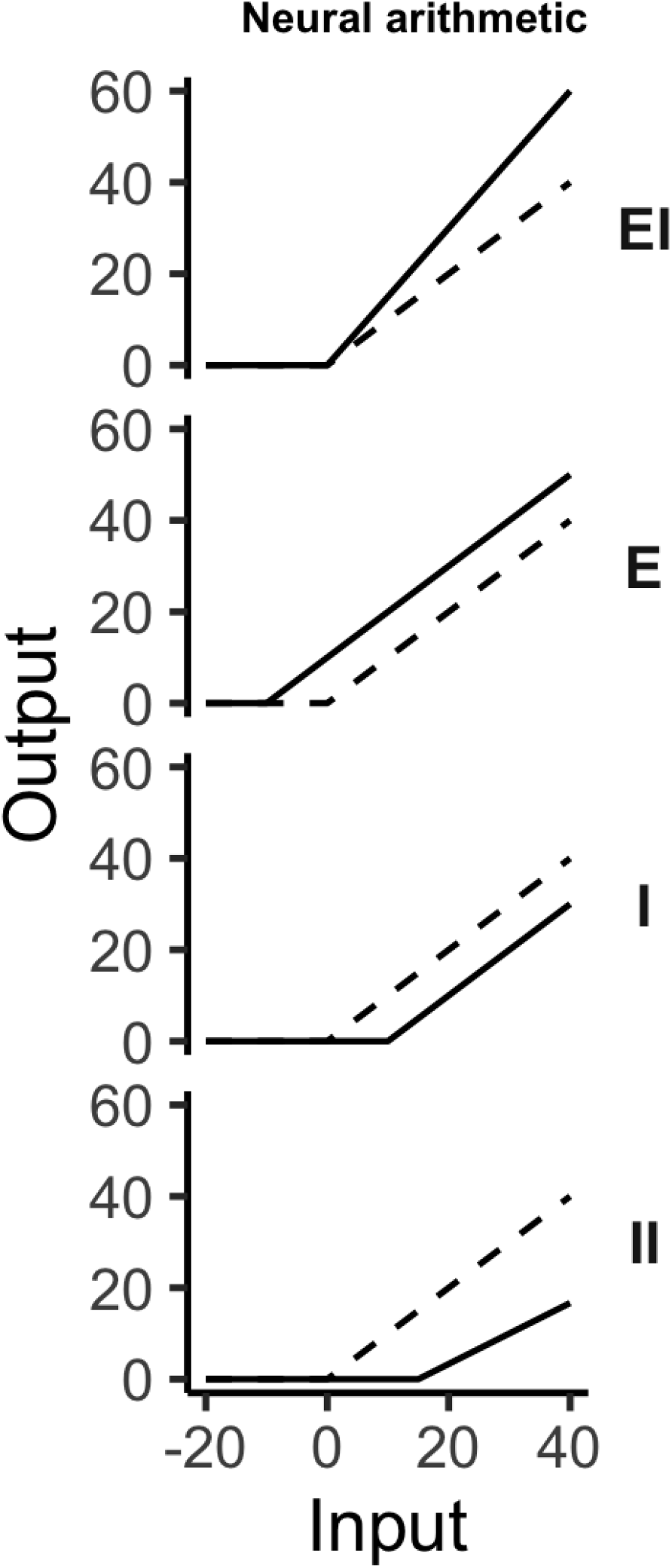
Neural arithmetic, as neural modulation.

Parameters and definitions for all equations, both the model and the metrics below, are summarized in Table 1

#### Biophysical circuit

Each biophysical model featured 2 or 3 populations. The total network consisted of *N* = 25000 Adaptive Exponential (AdEx) neurons (61) This total was subdivided into populations with distinct biophysical parameters (Table 2). These parameters were tuned to human recordings, as detailed in (33). Each population was connected to the others in a model-specific way (Figure 1). All shared the common stimulus drive (Eqs. 4 and 6).

Our biophysical model was based on open source code from (33), available at http://modeldb.yale.edu/267039. We now review this model, but include only the details necessary for a reproduction of our results. For justification of the default parameter, see (33).

All membrane dynamics were governed by an AdEx model (Eq. 13).

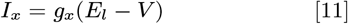

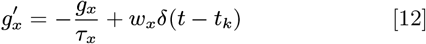

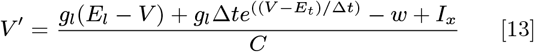

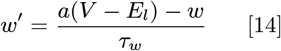

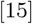

Here the leak conductance *g_l_*, the capacitance *C*, and the leak potential *E_l_* control the passive properties of each neuron.

While action potential initiation is driven by the exponential Δ*te*^(.)^ (Eq 13). Following an action potential, the membrane has both a “fast” step, where *V* and *w* are reset (Eq. 16), and a “slow” (passive) response governed by dynamics of *w* itself. Here *a* and *τ_w_* define the rate of change of *w* outside of “fast” events.

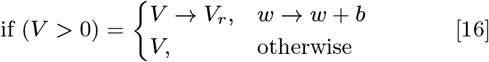

The parameters we explored to modulate oscillatory translation in our ING, PING and CHING models are shown in Table 3. The connection probability from the drive stimulus to all target populations was *p* = 0.02, and consisted of *n* = 2500 Poisson cells.

1. Model ING –Consisted of three populations. One set of 20000 Regular Spiking (RS), one set of 4000 Fast Spiking cells (FS1), and another small set of FS cells, labelled FS2 (1000). RS cells were excitatory (E). FS1/FS2 cells were inhibitory (I). The FS2 population was the driver of the gamma rhythm. It featured connection with a 0.6 probability, synaptic strengths of *g_I_I* = 5*nS* a *τ_I_* = 5*ms* time constant and a connection delay of 1.5 ms. FS2 cells have random recurrent connections with RS neurons with *p* = 0.15. FS2 cells have random recurrent connections to FS with *p* = 0.15, while FS cells send connections to FS2 neurons with a probability of *p* = 0.03. Synaptic strengths of all excitatory connections were *g_E_* = 1*nS*. All inhibitory connections were set to *g_I_* = 5*nS*. Synaptic half-lives and delay time were equal in this network, set at *τ_E_* = *τ_I_* = 5*ms* and 1.5 ms respectively. The drive stimulus was excitatory to all populations, *g_D_* = 0.9 * *nS*.
2. Model CHING – Consisted of three populations. One set of 20000 Regular Spiking (RS), one set of 4000 Fast Spiking cells (FS1), and another small population of 1000 excitatory Chattering cells (Ch). Connections between cells were made with a 0.02 probability. Synaptic half-lives and delay time were equal in this network, set at *τ_E_* = *τ_I_* = 5*ms* and 1.5 ms respectively. Synaptic strengths of all excitatory connections were *g_E_* = 1*nS*. Inhibitory connections from FS to RS and FS to Ch were set to *g_I_E* = 7*nS*, while internal inhibitory connections from FS to FS were set to *g_I_E* = 5*nS*. The drive stimulus was excitatory to all populations, with population specific synaptic strengths of, *g_DRS_* = 1.0, *g_DFS_* = 0.75, *g_DCh_* = 0.75*nS*.
3. Model PING – Consisted of two populations. One set of 20000 Regular Spiking (RS) and one set of 5000 Fast Spiking (FS) cells. RS cells were excitatory (E). FS cells were inhibitory (I). Connections between cells were made with a 0.02 probability. All excitatory connections had the same default conductance, *g_EI_* = *g_EE_* = 5*nS* and a 1.5 ms connection delay. As did all inhibitory (I/FS) cells, *g_IE_* = *g_II_* = 3.34*nS*, though these had a longer 7.5 ms delay. The drive stimulus was excitatory to all populations, *g_D_* = 0.8 * *nS*.

### Mutual information

To calculate entropy and mutual information we took a discretization approach, in particular a binning approach. This worked as follows. First all signals were normalized between [0, 1]. The [0, 1] range was then divided into *m* bins. We set the bin number *m* to be equal to the average stimulus drive, *r_s_*. That is 10 spks/s, which is the same as 1 Hz. See Figures 3-4 for an example. Our rationale was that 1 Hz is a reasonable first pass estimate for how precise a downstream decoding population can be. Doubling or halving *m* did not qualitatively change our findings (Figure 9).

After discretization we constructed histograms separately from the reference signal and target firing. Normalizing these histograms by the total count generated discrete probability distributions from which we can calculate entropy *H* and mutual information MI (Equations 17-18). Examples of these distributions can be found in Figure 4.

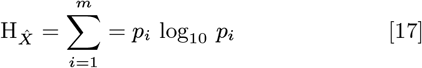

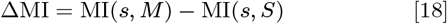

To measure translation cost/benefit we made two of these information theory calculations. We first measure the information shared between a time-varying stimulus *s* (whose creation is described in the *Methods*) and a Poissonic firing-rate sample of that stimulus 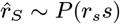, where *r_s_* is some average firing rate. That is, MI(*s, rs*). Second we measured the information shared between the stimulus *s* and a sample of the *modulated* stimulus. That is, 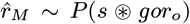 where *o* is the time-varying oscillator (see *Methods*), *g* represents the fixed “synaptic” strength of the modulation, *r_o_* is the average firing rate, and ⊛ represents one of the four modulations outlined above. This gives, 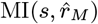. For notational purposes we replace the *M* the 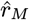 with the notation for modulation itself. For example, in model E we would denote the *M* to read as *r_E_*. The final cost/benefit measure is given by Eq. 1 (*Results*).

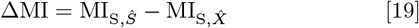

The central idea behind our use of ΔMI is that if oscillatory modulation is truly beneficial it should do better, that is have higher MI, than just sampling *r_s_* alone.

## Discussion

We have made a beginning of a theory to understand benefits and costs of oscillatory translation in a rate setting. We considered both a model of an independent oscillator, and three biophysical circuits tuned to match human electrophysiology. In both models we observed that oscillatory entrainment is beneficial overall, that is when shifting from an aperiodic code to a periodic one, when the oscillation can add new spikes bring the population and stimulus closer together. The ranges of when this can occur are highly circuit specific. To explain the benefits and limits of simulations, our discussion takes the form of questions and answers.

### So physiological details matter, a lot?

In both models the results we find depend on factors hidden in a typical electrophysiology experiment for studying oscillations–ie the LFP. The population size and synaptic weights, for example.

A debate which has been happening since oscillations were noted can be summarized by, “do oscillations do anything, or are they a byproduct of some other process?”. In some ways this question is settled. We have clear examples of oscillations seeming to cause neural pathology (62, 63). But for much of the literature the link to a direct functional role rests on correlative findings or interventions with weak effect sizes. So in some ways the basic question posed above is still an important one.

This basic question is why we are interested in costs. Oscillations arise naturally in most any system of equations which has additive, subtractive, or balanced interactions (10, 42, 58, 59, 64–66). It is natural to wonder if evolution has taken these general mathematical phenomena to harness. If so, then costs are a central issue because to take advantage of rhythms, their use would develop in accordance with their limits.

### Why would we use a rate model?

Why would we assume a rate model? Why not consider spike-timing or coincident codes?

The short answer is we have a separate line of work where we study oscillations compression algorithm (67) which *precisely* exchanges degrees of freedom in spike-timea for synchronization.

The other answer is that rate models are used to fit a very large range of electrophysiology data and in some cases to pre-test and validate clinical results. Epilepsy is a well known example (68), but there are others (69). Despite their simplicity then, or perhaps because of it, rate models used in the clinic and for other interventions require that they capture something real in the data (70). More than curve fitting, but in some fashion mechanistic.

But we argue one of the most interesting things about oscillations as a phenomenon is that they are ubiquitous across circuits, and even animal phyla. We hypothesize this ubiquity argues for simplicity in mechanism, ala rate codes. Even if we are wrong about oscillation’s mechanism as rooting in simplicity, and rate codes, then at the very least our simple view was worth considering first.

### Why be so opinionated about the code?

(33) asked a series of questions similar in spirit to ours, and in this work measure integration, coincidence detection and resonance in the same biophysical model we used. (We used their code as the basis for our experiments, in fact). They reported parameter regimes where gamma oscillations appear are least responsive, and have no more spike-time coincidences than aperiodic areas of parameter space.

Though they do not use a concept of cost/benefit in framing their results, all their results are consistent with gamma oscillations as a purely costly phenomena. At least in some looser sense,

The big difference between our two analyses is the assumptions we make. By presuming a rate code was can observe exact message fidelity, using information theory. This let us detect some benefits to gamma oscillations which were otherwise missed.

### Why do ING and I disagree?

The benefits of synchronizing firing in INg models was previously shown in (18), though their approach requires more specific wiring conditions than we enforce here.

### Are costs always costs? Are benefits always benefits?

The answer to these questions is a definitive no. The cost/benefit labels we have been using follow from our assumption that oscillations aid communications. We could have adopted the opposite assumption, which is just as supported by theory and experiments. That is, oscillations can gate out information, or otherwise suppress communications (71–73). So from this view what we have been calling a cost (*-ΔMI*) is not, but can be seen as an optimal result. And what we have been calling a benefit must be seen as a kind of cost, or error.

### Transformation, not transmission

We assumed oscillations are a communications channel. This view of oscillations is common today, so we believe we are justified in the work of others (40, 74). However from another view the “goal” of neurons is not broadly speaking to transmit information, but to compute with it (75–79). If computation is a rhythms’ primary role, then the analysis in this paper does not seem to apply and further work is needed.

One of the closest analyses to ours was Neymotin et al (50), who studied both translation and transformation of information flow in cortical microcircuits. They focused on the impact of recurrent connections as a source of past knowledge. These kinds of connections have a tendency to be beneficial, for the simple reason that past memories are often useful for future actions. However like in our work, when intrinsic connections grow to strong past compromises new information flow. Translation by oscillation is different in one important way. Oscillations can act as a “perturbation” not necessarily a memory. This perturbation can organize, gate, or amplify. But these same mechanisms allow it to also disturb functioning–as shown in (80, 81).

### What are the new predictions?

We assume in all cases the ability to make dual recordings of spiking at two locations in the central nervous system, and that the connection type(s) between these locations are known. For example, AMPAergic (E) or GABAergic (I) connections. With the ability to make dual recordings, any experimentalist can construct rate-based information theory measures, like those we studied in this paper.

Our predictions are qualitative, and seek to establish a basic concordance between our model and a real nervous system, to be measured. We assume also that passive recording of cortical areas can generate sufficient variability to test these predictions. We predict,

1. For an independent oscillation a rise and then decline in |Δ*MI*1 follows the size of the population from which it is later decoded. This is predicted to occur regardless of type of oscillatory entrainment in this model class.
2. For recurrent circuits who entrain via inhibitory interactions (ING or PING), |Δ*MI*1 increases monotonically with population size, up to at least 10000 neurons.
3. Direct (monosynaptic) inhibitory I oscillatory coding should uniformly show negative Δ*MI*, with the exception of large (>10,000) populations subjected to a strong driving stimulus.
4. Whereas for ING with full recurrent connections oscillatory coding should show positive Δ*MI*. This effect should increase further as the inhibitory conductance increases.
5. Direct (monosynaptic) excitatory E oscillatory coding should uniformly show positive Δ*MI*.
6. Whereas CHING entrainment in a recurrent circuit begins with small positive Δ*MI* that declines with population size, and shows little dependence on stimulus firing rate or conductance strength.
7. Oscillations that affect both E and I connections with an independent oscillation that are balanced (41), lead to qualitative distinct cost/benefits when compared to E or I direct connections. Namely,
8. Increases to the stimulus drive increase Δ*MI* for EI connections, but
9. Increases to the stimulus drive decrease Δ*MI* for E connections.
10. For large populations, increases to the amplitude of the oscillation will shift Δ*MI* from a positive value for in EI connections, to a negative value.
11. But this will not happen for E connections.
12. In contrast, all recurrent models show monotonic decreases in Δ*MI* as the stimulus firing rate rises.
13. In ING and PING circuits these stimulus driven negative Δ*MI* are mitigating by increasing oscillatory coupling conductances,
14. While the opposite pattern emerges in CHING, where weaker oscillations can mitigate stimulus drive decreases in Δ*MI*.

#### Summary

We have made the beginnings of a cost-benefit theory for oscillatory codes, and shown how the effect of oscillations can be quite intricate when measured in information theoretic terms. Both intrinsic connections, inputs, and population size can have about equal effects on costs. The details matter, in other words. A fully realized version of this theory would let us predict when and where oscillation would aid, or harm communication, *and by exactly how much*.

## Addendum

This work is supported by a Sloan Research Fellowship (FG-2015-66057), the Whitehall Foundation (2017-12-73), the National Science Foundation under grant BCS-1736028, and a Halicioğlu Data Science Institute Fellowship (to B.V.). The authors declare that they have no competing financial interests. Correspondence should be addressed to E.J.P.

